# Variation in cultural attitude, knowledge and individual motivational factors impact engagement and tool use in a field experiment in wild chimpanzees

**DOI:** 10.64898/2025.12.01.691567

**Authors:** Kelly Ray Mannion, Ndora Michael, Kugonza Stephen, Thibaud Gruber

## Abstract

Cultural traditions shape how animals approach and solve problems. Previously, Ugandan chimpanzees (*Pan trogodytes schweinfurthii*), have engaged with the honey-trap experiment, an apparatus designed to mimic a beehive and provide ecological opportunities for tool use, relying on their cultural knowledge. Here, we presented chimpanzees from the Mwera South community, a newly habituated community in Bugoma Forest, with variations of the honey-trap experiment and compared their engagement to other Ugandan chimpanzee communities to investigate various aspects of cultural behaviour in animals. First, we wanted to test whether communities varied in cultural attitude towards a given food, and whether this attitude correlated with particular instrumental components of their cultural knowledge. Second, we were interested in analyzing individual variation across individuals within the same culture. Comparing individuals from Bugoma’s Mwera South (N=15), Budongo’s Sonso (N=34), and Kibale’s Kanyawara (N=14) communities, we found that the latter both exhibited higher engagement with the honey-trap, and a dedicated instrumental method, stick use, to obtain the honey. In contrast, Sonso and Mwera South chimpanzees appeared similar in lacking a cultural attitude towards honey, with no attached tool technique overall. Nevertheless, there were also strong inter-individual differences. Notably, some Mwera South chimpanzees displayed undescribed behavioural flexibility, using a range of tool behaviours including both stick and leaf tool use —a pattern never before documented in over a decade of honey-trap experiments. These results demonstrate that cultural attitudes toward resources constitute an additional layer to culture in addition to instrumental knowledge, and that the latter two coexist with individual motivational traits in influencing the realization of cultural behaviour.

## Introduction

Decades of research in established long-term field sites yield a plethora of evidence in support of a growing number of animal species having culture. In particular, over the last two decades, evidence for social learning has been found across many taxa, whether in avian, mammalian or even in insect species, and in a range of behaviour such as foraging or vocalizations [1–5]. Specifically, these cultural species have displayed the potential and ability for social learning that enables the transmission of behavioural traditions that are passed on through generations [6–8]. Correspondingly, research has also looked into whether, as in humans, and how an individual’s culture influences their behaviour in nature or in the context of field experiments [9].

This cognitive approach to culture has been complemented in recent years by wider approaches to what we can consider cultural in animals. For example, Schuppli and van Schaik [10] argue that by focusing only on remarkable behaviour such as tool use, we “underestimate culturally transmitted basic subsistence skills” (p.3) such as revealed in diet [11]. Others have argued for differences in what can be characterized as *social attitudes* to count as cultural variants [e.g. 12, 13]. The two approaches are not necessarily incompatible. For example, a particular attitude towards a specific food resource may either favor or disfavor its consumption inside the population. Many examples are found in humans (for example, many English people are fond of marmite, while most others, including in the UK, are fairly unmoved by its taste). This culturally-acquired taste for a given food may drive attitudes towards this particular food, as well as instrumental learning, including tool use, to access it.

The latter point illustrates a key aspect of cultural knowledge: while a general attitude towards a given object may be socially acquired – one may here talk about the acquisition of *value*, see [14, 15] – how individuals within a community will implement this cultural attitude will also possibly vary, particularly if a given instrumental action is connected to it. For example, if a particular resource is highly valued in the community, and there is a dedicated technique to access it, one can predict that this technique will prevail in the community. To the contrary, if there is only a weak interest for a given object or resource, there may not be a set technique dedicated to obtaining the latter. As such, we can predict that communities where there is only a weak interest for a given resource will not have a set attached behaviour towards the food reward. This, however, does not exclude individual preferences, and such, does not preclude some individuals from the community from having a certain knack for the object of interest and making use of their existing knowledge, cultural or not, to obtain that resource.

To test hypotheses in the field rather than a laboratory or captive context, researchers have implemented field experiments on wild populations, which deliver more ecological validity compared to semi-wild or captive contexts. Field experiments testing foraging behaviours can be used to examine cultural tendencies such as tool use in foraging scenarios, for example by presenting a substrate and the required tool material [16, 17]. For example, Gruber et al. [18, 19] developed the honey-trap experiment, which simulates a natural context, such as a beehive, where the honey is only accessible through a small hole, thus presenting the opportunity for the chimpanzees to use a tool to extract the honey. Eastern African chimpanzee (*Pan troglodytes schweinfurthii*) communities in Ugandan Budongo and Kibale forests displayed community specific responses to extract honey from the apparatuses, relying on their already established foraging tool use behaviour. Crucially, they did not develop behaviour outside of their cultural knowledge. More recently, Koops et al. [20] replicated these results in another subspecies, Western African chimpanzees (*P. t. verus*), and with a different tool behaviour: nut-cracking using stones. Combined, the results of these studies suggest that foraging field experiments have little impact on triggering or modifying wild chimpanzee behaviour, beyond multiplying opportunities for chimpanzees to express somewhat rare behaviour without modifying their cultures, an ethical issue [21]. As such, they can rather be seen as a revelator of the current cultural behaviour of chimpanzees. In addition, by solely providing opportunities for engagement, field experiments may also act as a proxy for general interest in the resource. For example, the Kanyawara chimpanzees of Kibale Forest appeared to display general differences in their interest in honey compared to the Sonso chimpanzees of Budongo Forest [22].

Ugandan chimpanzee populations appear particularly suited for testing questions related to cultural attitudes and individual differences regarding food items. Cultural repertoires, including tool use behaviour, have been described for several Ugandan chimpanzee communities due to long-term field sites with research and conservation projects. The innovation and subsequent transmission of moss-sponging to acquire water has been well documented in the Sonso community of Budongo Forest [2, 23]. However, unlike most other chimpanzee communities, the Budongo chimpanzees have not been recorded using stick tools (for a review see [22]). Stick use is however described throughout Uganda including in Kibale Forest [24], Kalinzu Forest [25] and in neighboring forest patches to Budongo such as Kasongoire or Bulindi [26]. The distinction in stick tool use between Kibale and Bulindi chimpanzees is also of interest with respect to cultural knowledge: while Kibale chimpanzees use stick tools to probe, Bulindi chimpanzees use stick tools to dig in the ground and unearth beehives [27].

The mosaic of chimpanzee communities in Uganda makes studying the variation in tool use and thus chimpanzee culture well-grounded. The Bugoma Forest lies in the middle between Kibale and Budongo forests, with the resident chimpanzee communities exhibiting marked cultural differences with other chimpanzee communities such as their nest building behaviour [28, 29]. Yet, their foraging cultural toolkit remains elusive, with possible observations of stick use being suggestive of a stick tool culture (see Supplemental Figure 2). Here, we presented the resident Mwera South chimpanzees of Bugoma Forest with variations of the honey-trap experiment to allow them to express their foraging strategies in a controlled environment. We replicated two previous studies with the Budongo and Kibale chimpanzees to offer them the same ecological affordances as previously offered to the other two communities. In particular, we both presented the standard honey-trap experiment (a hole carved into a large log, [18]), and the same setting complemented with offering a leafy-stick tool [19]. The addition of this third chimpanzee community allows triangulating the aspects of interest, attitude and instrumental means towards a resource as described above. We first compare the results of the experiment across the three communities and show that the Kanyawara chimpanzees appear as an outlier in their demonstrated interest for honey compared to the Sonso and Mwera South chimpanzees. This suggests that there are community-level differences in terms of attitude towards a given resource. Nevertheless, in all communities we find outliers who appear to particularly enjoy the resource. These chimpanzees are also the ones who use the largest diversity of means to access honey. In particular, we describe flexibility in tool using in individuals from the Mwera South community, including one who displays both stick manufacturing and leaf-sponging to solve the problem, raising questions about the affordances of the task. We discuss the interaction between community-specific factors and individual factors such as motivation in chimpanzees expressing their tool-using behaviour.

## Methods

### Study Site and Subjects

Bugoma Central Forest Reserve, Uganda (01°15′N 30°58′E) is a semi-deciduous tropical rain forest located between the Central Forest Reserve of Budongo (425□km2) and Kibale National Park (776□km2). The Bugoma Primate Conservation Project (BPCP, www.bugomaprimates.com) has been studying the Mwera South community since 2016 with habituation and identification of individuals in the community still ongoing. Currently, the community is estimated to have about 70 individuals similar to other East African chimpanzee community sizes. Research efforts with the Mwera South chimpanzees have already established the use of leaf-sponging on a usual basis to obtain water and leaf napkins/wipes (see Supplementary videos) as well as the existence of specific cultural behaviour such as ground nesting [28]. Because habituation was ongoing at the time of the study, it was unknown prior to the experiments what other tool use behaviour the Bugoma chimpanzees had in their repertoire.

The Sonso community (01°43’N, 31°32’E) inhabits the Budongo Forest Reserve, a 482 km² area of continuous medium-altitude semi-deciduous forest at approximately 1050 m elevation. Habituation of the Sonso community began in 1990 and was considered complete by the time of the experiments. The Kanyawara community (00°33’N, 30°21’E), located approximately 180 km from Sonso in Kibale National Park, has been continuously studied since 1987 and was fully habituated by the time of the experiments. Both communities practice leaf-sponging. Notably, Sonso chimpanzees do not display any other foraging-related tool use [2]. Kanyawara chimpanzees also exhibit fluid-dip behaviour habitually. Data in the honey-trap experiment for these communities came from studies in Gruber et al. 2009 and 2011.

### Experimental set-up

The set-up is only described for Bugoma in the current study. For data in Budongo and Kibale, please refer to Gruber et al. [18, 19] and Gruber [22].

Logs from fallen trees were collected from the forest and used to create experiment apparatuses. We cut a rectangular-shaped hole (4cm x 5cm x 16cm) into them using a hammer and chisel. After creating the hole, experiment logs were always disinfected and left to rest in a nearby clearing prior to being set up in the forest. Any personnel who handled the experiment apparatus or equipment for set up was required to wear face masks and gloves to avoid contamination (see [21] for ethical aspects of field experiments including risks for disease transmission).

The prepared logs served as the basis of the honey-trap experiment. Honey was poured into the hole of the log once set up in the forest until there was a 10cm gap from the top of the honey in the hole to the surface of the log, replicating Gruber et al. 2009’s setting (obligatory condition). Afterwards, pieces of honeycomb were placed on top of the hole (Figure 1) and held in place by beeswax to help providing a visual cue to the chimpanzees as well as deterring bees and other insects from accessing the honey. There were no additional means used to attract individuals or encourage them to engage with the experiment. Honey, honeycombs and beeswax for the experiments were harvested from hives maintained by local beekeepers in the villages surrounding Bugoma Forest Reserve. The bees were of the genus *Apis* and could move freely within and around the forest area.

**Figure 1:**
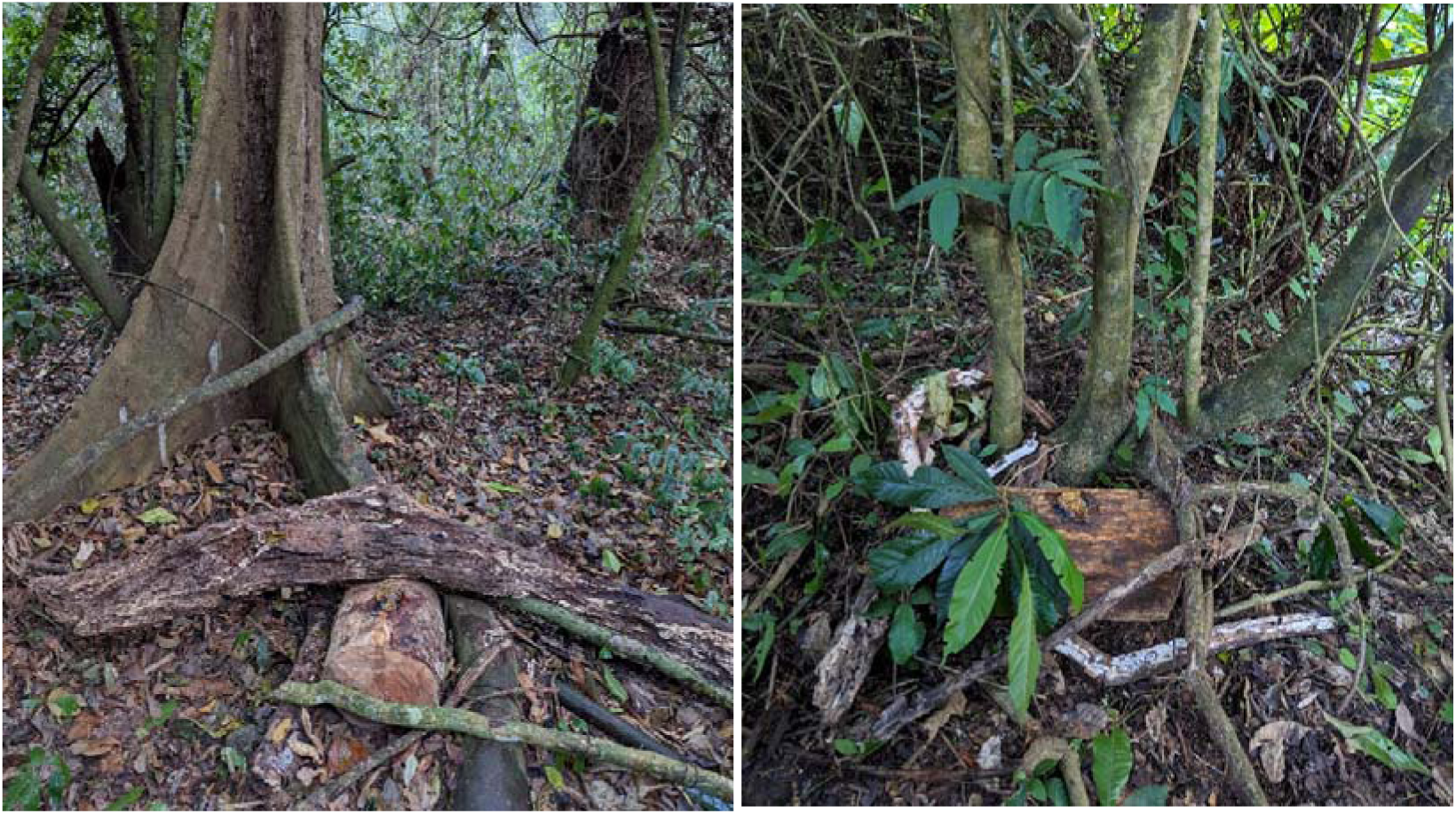
(left) Honey-trap Experiment 1. Honeycomb held in place over the hole with beeswax. The experiment is fixed to the tree buttress. (right) Honey-trap Experiment 2 with a leafy stick tool provided.

Experiment locations were chosen based on chimpanzee activity at the time of the experiment setting. Trails and resting areas near feeding trees were used depending on the frequency upon which the chimpanzees were using these areas during their normal day-to-day routine. Experiment areas preserved the normal forest underbrush and leaf litter present ensuring the most naturally simulated context and providing potential raw materials for the chimpanzees to use.

Multiple motion sensing camera traps (Bushnell CoreDS) were set up around the experiment, ensuring the visibility of the experiment log and unobstructed angles of individuals that engaged with it. Each experiment was set up prior to the chimpanzees arriving in the area to ensure that researchers would not be associated with the experiment. After setting up the experiment, researchers left the area and avoided any interaction with the chimpanzees.

Experiment 1 consisted of the honey-trap experiment in the obligatory condition. Experiment 2 consisted in the honey-trap in the obligatory condition experiment with the addition of a leafy stick laid on the top of the experiment log between the hole and the log edge to serve as a potential tool with multiple functions (see [19]). The side of the experiment log which the tool was placed on changed randomly with the experiment trials. The potential leafy stick tool was a 40cm branch/sapling of *Alstonia sp*., which is commonly found in Bugoma Forest. The lower 20cm portion of the tool had the leaves removed so that the tool could be used in different ways: to dip into the apparatus using just the stick end without leaves, to leaf-sponge by removing leaves to chew into a clump to dip into the hole and extract the honey like using a sponge, or as a combined leaf-dipping tool by inserting the leafy end of the stick and sucking honey off of the attached leaves that were dipped into the hole. The latter behaviour was never displayed by either the Kanyawara or Sonso chimpanzees (see [22] for a review). This experiment set-up tests whether chimpanzees can identify a potentially useful tool to extract the honey.

Between May 2021 and November 2022, we focused on Experiment 1, while in 2024, we focused on Experiment 2. The research was reviewed and received approval by the Uganda Wildlife Authority (permit n° COD-96-05 to KRM) and the Uganda National Council of Science and Technology (permit n° NS155ES to KRM).

### Video data in Bugoma Forest

Data were extracted from the videos using BORIS software [30]. Identities of chimpanzees and classification of behaviours were coded by creating a customized ethogram specific for behaviours concerning wild chimpanzees in a foraging context. Data recorded from the videos were exported from BORIS as a .csv file and imported into Rstudio for further processing and analyses.

### Data analysis

Data for Bugoma were directly coded from the video data by KRM. Data for the honey-trap experiment from the Sonso community in Budongo and Kanyawara community in Kibale were extracted from Gruber et al. 2011 (Table 1, “Baseline” experimental condition with non-zero total interaction time) and compared to the data acquired from the Mwera South community in Bugoma in the present study. All data duration in Mwera South were coded to the ms on BORIS and analyzed as such in statistical analyses, but rounded to the seconds in tables and figures for comparison purpose. Overall, we present data for *N_Sonso_* = 34; *N_Kanyawara_* = 14; *N_Mwera-South_* = 15 chimpanzees. An outlier was considered as an individual whose engagement time was more than 3 standard deviations from the overall mean across all communities (pooled threshold). This identified two individuals: QT from Kanyawara (z-score = 5.9) and HON from Mwera South (z-score = 3.6), which were subsequently removed for statistical analysis.

**Table 1:** Summary statistics of comparison between Kanyawara, Mwera South and Sonso during the baseline condition, Experiment 1.

| *Comparison* | *Dunn's Z* | *Dunn's $p_{adj}$* | *MW r* | *MW $p_{adj}$* | *Median diff. (s)* |
| --- | --- | --- | --- | --- | --- |
| *Kanyawara vs Mwera* | 2.49 | 0.019* | 0.438 | 0.092 | 286 |
| *Kanyawara vs*<br>*Sonso* | 2.22 | 0.04 | 0.454 | 0.044* | 219 |
| *Sonso vs Mwera* | 0.708 | 0.719 | 0.169 | 0.356 | 67 |

### Statistics

We first compared total community engagement using the same standard as in Gruber et al. 2011 to allow for comparison. However, unlike Gruber et al. 2011, where individuals who interacted for less than 20 s were discarded, we used all individuals unless they were statistical outliers (engagement durations > 3 SD from the mean). To test for community-level differences, we used a Kruskal-Wallis test with Dunn’s post-hoc pairwise comparisons (Bonferroni correction) and Mann-Whitney pairwise tests (Holm correction), both non-parametric approaches appropriate for skewed distributions. As only one chimpanzee in Mwera South directly engaged with Experiment 2, we only compared the data for Experiment 1 for the Mwera South chimpanzees with the data of Table 1 in Gruber et al. 2011.

To compare latency before first tool use, we used a Kruskal-Wallis test with Mann Whitney pairwise tests (Bonferroni correction). For this analysis, we used data from Gruber et al. 2009 for the Sonso and Kanyawara chimpanzees. This analysis should be interpreted with caution given the small sample sizes, particularly for Mwera South (n = 3) and Sonso (n = 3) compared to Kanyawara (n = 11). Nevertheless, it is included here to highlight the large inter-individual variation found both within and between community. All analyses and raw data can be found at the following link: https://github.com/ChimpanzeeResearch/bugoma_tool_use/tree/main.

## Results

### Comparison of engagement in the honey-trap experiment across field sites

We conducted a total of 9 baseline experiments in Bugoma Forest with no tool provided, i.e. Experiment 1, which led to 2:55:22 (hours : minutes : seconds) of interaction by more than 16 individuals across 30 days (from a total of 46 days that had some presence of chimpanzees at the experiment). Similar to the Sonso chimpanzees of Budongo Forest, most individuals interacted by trying to use their fingers to reach the honey in the bottom of the hole or by biting or moving the experiment apparatus with brute force to get the honey. Of those that interacted with the experiment, only three individuals used a tool to successfully extract the honey (see below for a more specific description of Mwera South chimpanzee behaviour).

We found marginally significant differences in total direct engagement across the three communities (Kruskal-Wallis test: χ²(2) = 6.99, p = 0.03; Figure 2). Kanyawara chimpanzees (N = 14, median = 319 s, range: 44-2899 s) spent more time in direct engagement compared to both Mwera South (N = 15, median = 33 s, range: 2-1678 s) and Sonso chimpanzees (N = 34, median = 83 s, range: 4-493 s).

**Figure 2:**
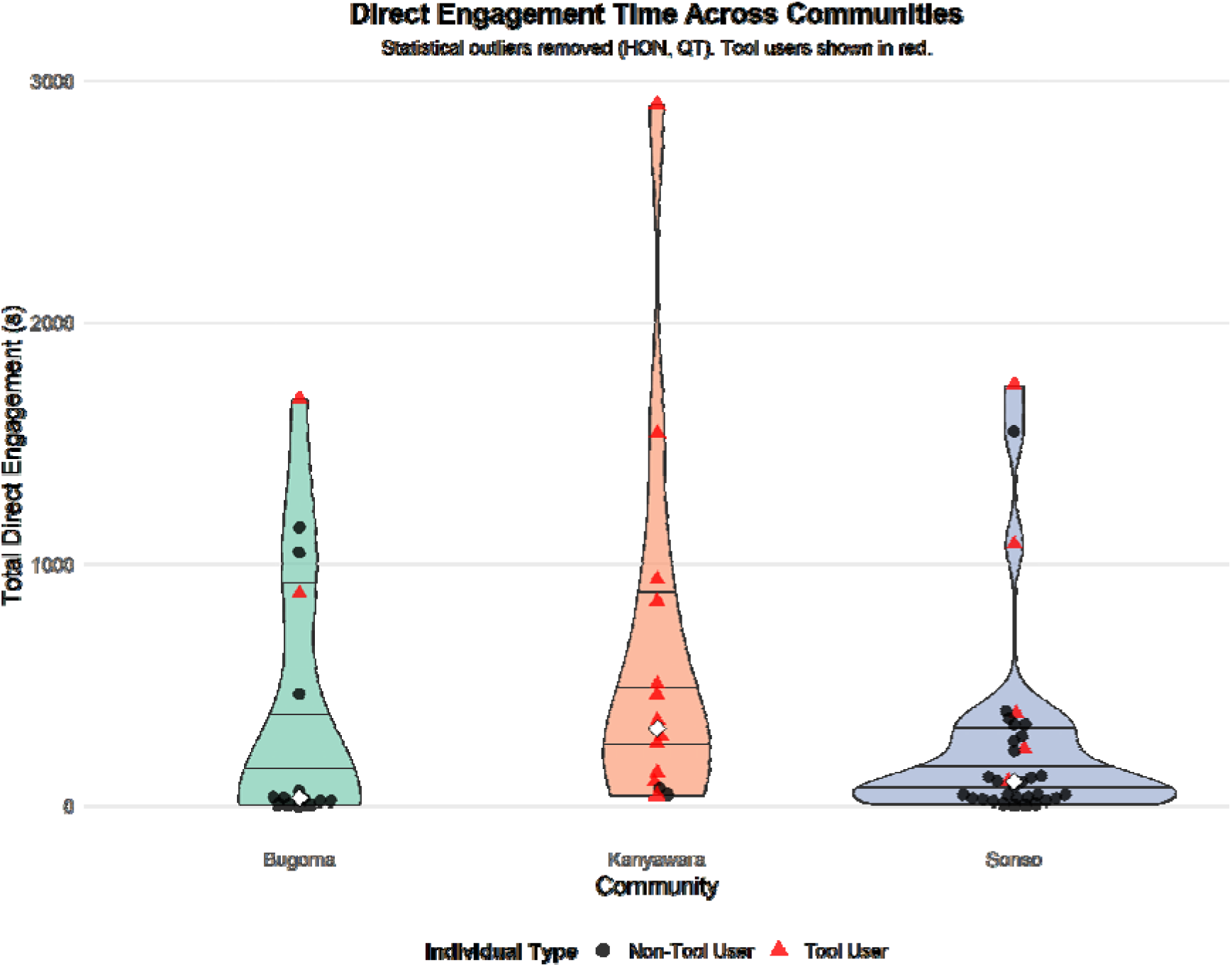
Distribution of total direct engagement time (in s) across three chimpanzee communities. Violin plots show the probability density of the data at different values, with individual data points overlaid (black dots for non tool users and red triangles for tool users). White diamonds indicate the median of each group. Note that Kanyawara shows the highest median engagement time (319 s; *N_Kanyawara_* = 14), followed by Sonso (100 s; *N_Sonso_* = 34) and Mwera South (32.6 s; *N_Mwera_* = 15), though with considerable overlap in their distributions. The width of each violin represents the kernel density estimation of the underlying distribution. Data for Kanyawara and Sonso is taken from Gruber et al. 2011. Both statistical outliers in Mwera South and Kanyawara are tool users.

Pairwise comparisons yielded different results depending on the test used (Table 1). Dunn’s test with Bonferroni correction (threshold p = 0.025) found that Kanyawara chimpanzees engaged significantly more than Mwera South (Z = 2.49, p = 0.019) but not Sonso (Z = 2.22, p = 0.040). In contrast, Mann-Whitney tests with Holm’s correction found a significant difference between Kanyawara and Sonso (r = 0.454, p = 0.044) but not Kanyawara-Mwera South (r = 0.438, p = 0.092). Both methods agree that there is no difference between Sonso and Mwera South.

In addition, there are noticeable qualitative differences between the communities: 13 of 15 Kanyawara chimpanzees used a stick in the experiments (86.7%), with none using a different tool, compared to five Sonso chimpanzees of 29 (17.2%) manufacturing a leaf-sponge (none used a stick). Finally, three Mwera South chimpanzees of 16 (18.8%; or 3 of 10, 30%, when only individuals above 20s are considered) used a variety of tool strategies (see below).

Latency to first tool use differed significantly among the three communities (Kruskal-Wallis test: χ²(2) = 6.71, p = 0.035). We observed considerable variation within communities, with Bugoma showing the highest variability (range: 55-1093 s, median = 111 s), followed by Kanyawara (range: 0-88 s, median = 20 s), and Sonso (range: 4-61 s, median = 54 s). Pairwise comparisons revealed that chimpanzees in Mwera South showed significantly longer latencies to first tool use compared to Kanyawara (Wilcoxon rank-sum test: W = 32, p = 0.057 after Bonferroni correction). The differences between Mwera South and Sonso (Wilcoxon rank-sum test: W = 8, p = 0.60 after Bonferroni correction) and between Sonso and Kanyawara (Wilcoxon rank-sum test: W = 9.5, p = 0.93 after Bonferroni correction) were not significant.

### Inter-individual variation

As shown in Table 2 and Figure 3, there was considerable within-community variation in engagement times, particularly in Kanyawara and Mwera South, while Sonso showed a more compressed distribution with 75% of values falling below 105 seconds. Kanyawara chimpanzees, while generally relying on the same tool-using technique, showed the largest variation in engagement time. The Mwera South chimpanzees are described below in more details.

**Figure 3:**
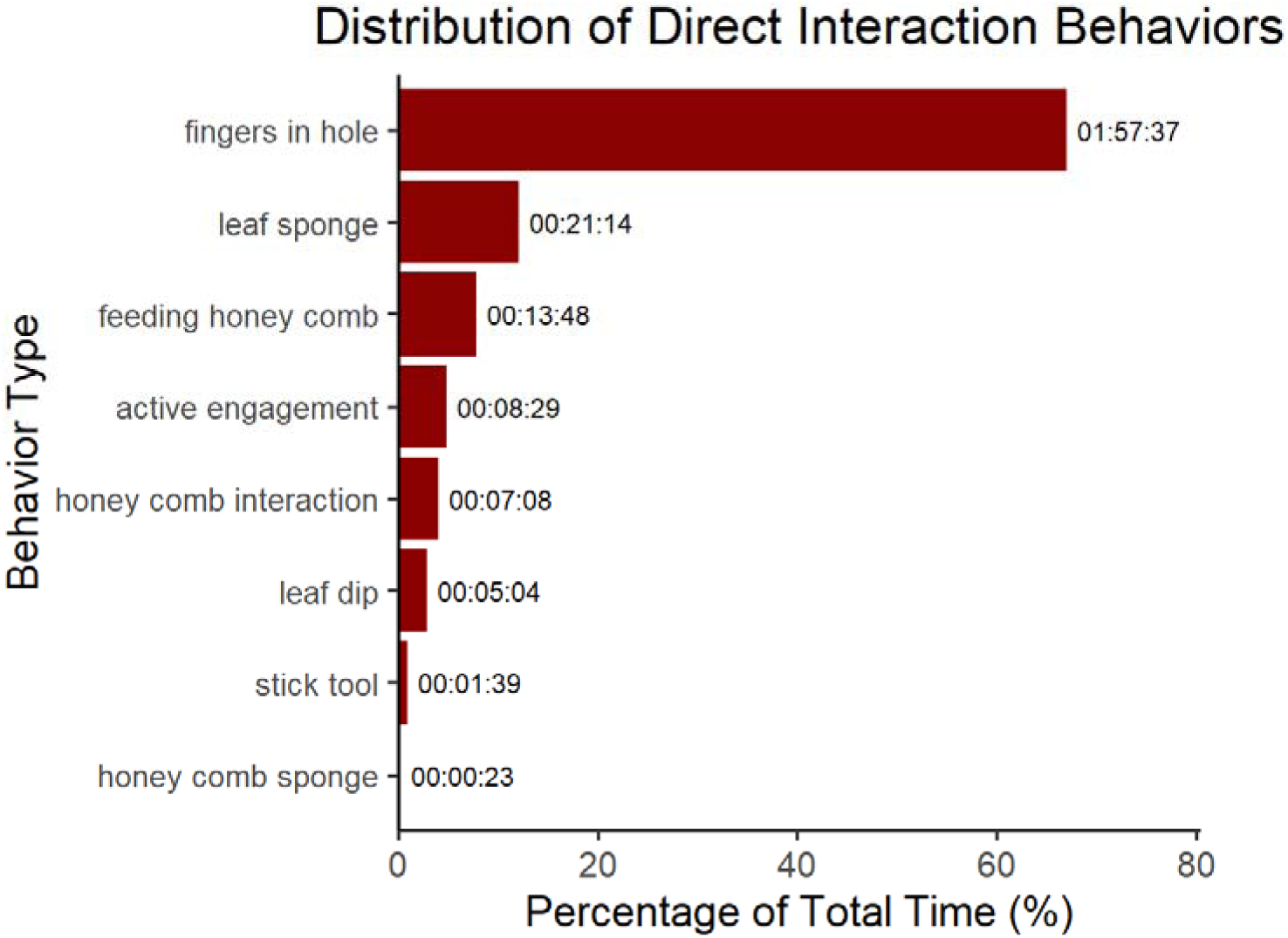
The frequency of direct interaction behaviour from the chimpanzees who engaged with the honey-trap experiment.

**Table 2:** Summary of individual engagement from chimpanzees during Experiment 1 of the honey-trap experiment across three Ugandan communities. *: individuals removed from the community-wide comparison because they were outliers statistically. Engagement time shown has been rounded to the nearest integer.

| **Community** | **ID** | **Age Class** | **Direct Engagement (s)** | **Tool** | **# Encounters** |
| --- | --- | --- | --- | --- | --- |
| Kanyawara | AL | Adult | 138 | Stick | 3 |
| Kanyawara | AT | Juvenile | 259 | Stick | 2 |
| Kanyawara | AZ | Juvenile | 460 | Stick | 1 |
| Kanyawara | BO | Juvenile | 97 | Stick | 1 |
| Kanyawara | ES | Sub-Adult | 286 | Stick | 1 |
| Kanyawara | EU | Juvenile | 846 | Stick | 2 |
| Kanyawara | KK | Adult | 72 |  | 1 |
| Kanyawara | LR | Adult | 502 | Stick | 1 |
| Kanyawara | NP | Sub-Adult | 2899 | Stick | 4 |
| Kanyawara | OG | Juvenile | 935 | Stick | 2 |
| Kanyawara | OT | Sub-Adult | 352 | Stick | 4 |
| Kanyawara | PG | Adult | 49 |  | 3 |
| Kanyawara | QT* | Adult | 7280* | Stick | 11 |
| Kanyawara | TJ | Adult | 1540 | Stick | 1 |
| Kanyawara | TT | Juvenile | 44 | Stick | 1 |
| Sonso | AN | Sub-Adult | 30 |  | 1 |
| Sonso | FD | Sub-Adult | 5 |  | 1 |
| Sonso | FK | Juvenile | 10 |  | 1 |
| Sonso | HL | Sub-Adult | 98 |  | 1 |
| Sonso | HT | Adult | 102 | Leaves | 1 |
| Sonso | HW | Adult | 381 | Leaves | 5 |
| Sonso | JT | Juvenile | 121 |  | 1 |
| Sonso | KA | Juvenile | 20 |  | 1 |
| Sonso | KE | Sub-Adult | 10 |  | 1 |
| Sonso | KL | Adult | 5 |  | 1 |
| Sonso | KR | Juvenile | 30 |  | 1 |
| Sonso | KS | Juvenile | 235 | Leaves | 1 |
| Sonso | KW | Adult | 78 |  | 3 |
| Sonso | KY | Adult | 335 |  | 1 |
| Sonso | KZ | Adult | 225 |  | 3 |
| Sonso | ML | Adult | 45 |  | 1 |
| Sonso | MN | Juvenile | 45 |  | 1 |
| Sonso | MS | Adult | 335 |  | 1 |
| Sonso | NB | Adult | 1547 |  | 4 |
| Sonso | NR | Sub-Adult | 391 |  | 5 |
| Sonso | NT | Juvenile | 1738 | Leaves | 5 |
| Sonso | RS | Sub-Adult | 1082 | Leaves | 3 |
| Sonso | SB | Adult | 360 |  | 3 |
| Sonso | SE | Adult | 45 |  | 1 |
| Sonso | SM | Adult | 45 |  | 1 |
| Sonso | SQ | Adult | 34 |  | 3 |
| Sonso | TK | Adult | 267 |  | 2 |
| Sonso | VR | Sub-Adult | 115 |  | 1 |
| Sonso | WL | Adult | 102 |  | 1 |
| Sonso | ZD | Juvenile | 33 |  | 1 |
| Sonso | ZF | Adult | 18 |  | 1 |
| Sonso | ZG | Sub-Adult | 119 |  | 3 |
| Sonso | ZL | Adult | 287 |  | 5 |
| Sonso | ZM | Adult | 25 |  | 2 |
| Mwera South | HON* | Adult | 4640* | Leaves | 13 |
| Mwera South | ENG | Adult | 1679 | Stick & Leaves | 5 |
| Mwera South | BRO | Adult | 1151 |  | 3 |
| Mwera South | KBY | Adult | 1048 |  | 3 |
| Mwera South | MAM | Adult | 876 | Leaves | 9 |
| Mwera South | ARP | Adult | 462 |  | 4 |
| Mwera South | JUV | Juvenile | 60 |  | 10 |
| Mwera South | AFO | Adult | 34 |  | 4 |
| Mwera South | SAM | Sub-Adult | 33 |  | 3 |
| Mwera South | CAP | Adult | 29 |  | 5 |
| Mwera South | AM2 | Adult | 18 |  | 5 |
| Mwera South | KAT | Juvenile | 18 |  | 3 |
| Mwera South | MKU | Adult | 7 |  | 3 |
| Mwera South | GER | Adult | 7 |  | 2 |
| Mwera South | AF2 | Adult | 4 |  | 6 |
| Mwera South | JUVF | Juvenile | 2 |  | 2 |

When directly engaging with the experiment, Mwera South chimpanzees relied on tool use 16.16% of their engagement time (see Figure 3 for exact breakdown). When specifically looking at honey extraction behaviours (see Table 3), chimpanzees relied on tool use 19.41% of the time and tried using their fingers the other 80.59% of the time (see Figure 3 for breakdown and Table 3 for behaviours). Interestingly, we recorded four separate types of tool-using behaviours by three individuals: stick tool, leafy-stick dip, leaf sponge, and honeycomb sponge. See Table 4 for a daily breakdown of tool use.

**Table 3:**
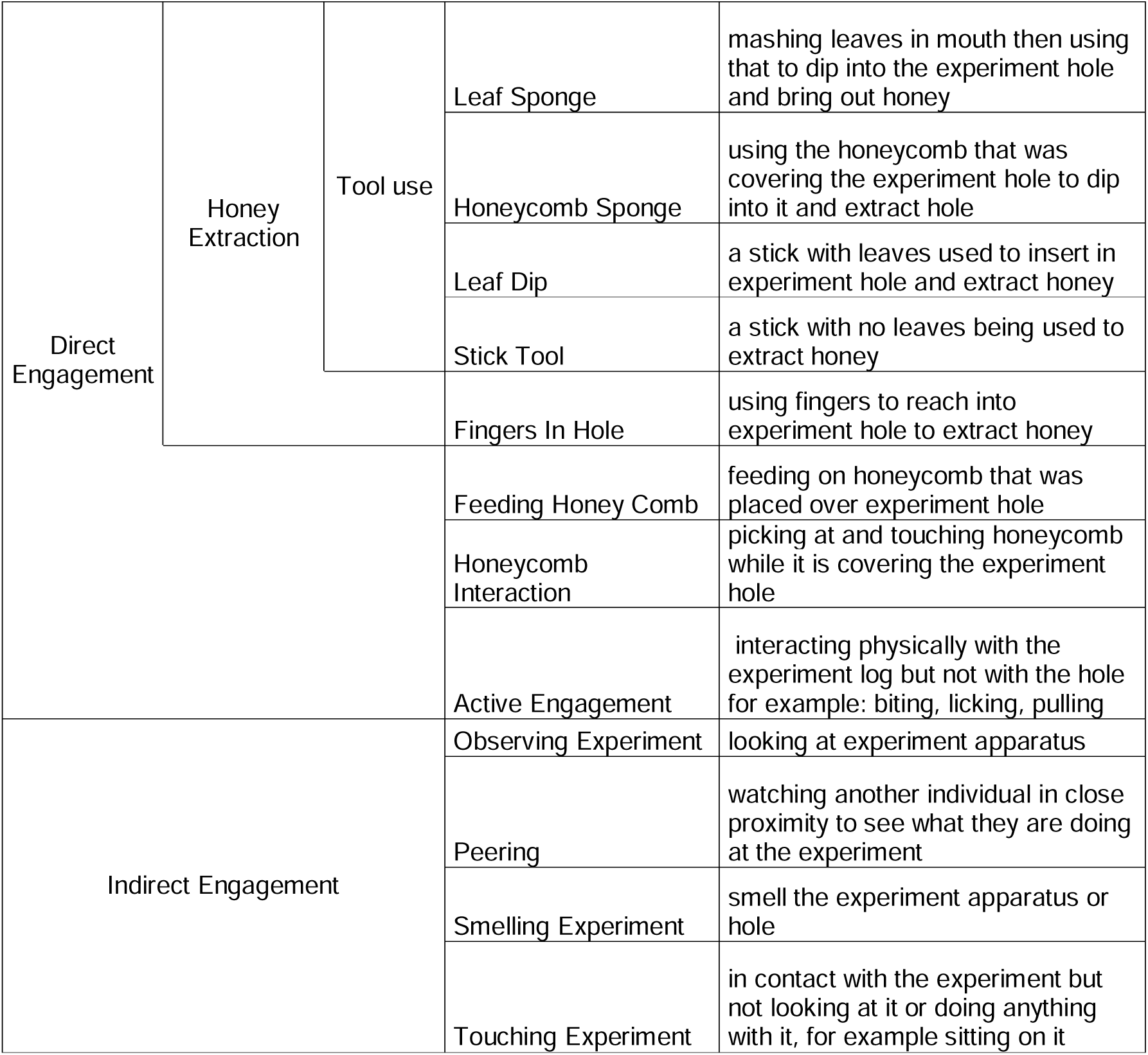
The description of different behaviour that the Mwera South chimpanzees exhibited during honey-extraction and the more general groups they were classified in.

**Table 4:** Latency to tool use by individual and experiment type for both experiment types. For each individual, the encounter column shows the day when that individual encountered the experiment, e.g. HON started using a leaf sponge the 2nd day when she encountered experiment with no tool provided. All times are in hours:minutes:seconds.

| Experiment Type | Subject | Encounter | Total presence that day | Latency to tool use | Direct engagement before tool use | Indirect engagement before tool use | Tool type |
| --- | --- | --- | --- | --- | --- | --- | --- |
| Notool provided | ENG | 1 | 00:29:55 | 00:21:40 | 00:18:12 | 00:00:53 | leaf dip, stick tool |
|  | HON | 2 | 00:15:07 | 00:05:13 | 00:01:50 | 00:01:27 | leaf sponge |
|  |  | 5 | 00:29:15 | 00:16:10 | 00:12:17 | 00:00:11 | leaf sponge |
|  |  | 6 | 00:17:53 | 00:06:55 | 00:06:55 | 00:00:00 | leaf sponge |
|  |  | 7 | 00:03:02 | 00:00:47 | 00:00:05 | 00:00:00 | leaf sponge |
|  |  | 8 | 00:14:59 | 00:00:36 | 00:00:31 | 00:00:03 | leaf sponge |
|  |  | 9 | 00:04:12 | 00:00:00 | 00:00:00 | 00:00:00 | leaf sponge |
|  | MAM | 2 | 00:11:26 | 00:02:14 | 00:00:55 | 00:00:30 | honey comb sponge, leaf sponge |
| Tool provided | ENG | 1 | 00:23:22 | 00:01:14 | 00:01:00 | 00:00:01 | leaf dip, leaf sponge, stick tool |
|  |  | 2 | 00:00:59 | 00:00:32 | 00:00:19 | 00:00:07 | leaf dip |
|  |  | 3 | 01:01:44 | 00:03:55 | 00:03:42 | 00:00:00 | leaf dip, leaf sponge, stick tool |

We now describe in more details the strategies used by the three tool-using Mwera South chimpanzees. A summary is provided in Table 4.

#### Leaf tools

As all other chimpanzees in Uganda, the Mwera South chimpanzees use customarily a range of leaf tools. In particular, the Mwera South chimpanzees regularly use leaf sponges to obtain water from holes and puddles where it collects after rain; they also use leaves as napkins to wipe themselves (see Supplementary videos). Experiment 1 yielded two female individuals, HON and MAM, who manufactured and used natural sponge tools to reach the honey in the experimental log. Experiment 2 yielded one male individual, ENG, who used a sponge tool.

#### Honey-comb tool

In the same manner as a leaf sponge, one of the females, MAM, first used the honeycomb as a sponge to get the honey from the experiment log, which has never been seen before in the wild.

#### Stick tools

An adult male, ENG, manufactured a stick tool to extract honey from the experiment log. ENG used both a leafy stick to dip into the experiment hole and get honey by sucking the leaves on the end of the stick (brush tool), as well as manipulated the stick to make a bare stick with no leaves on it to dip into the hole and retrieve the honey (Figure 4, see Supplementary videos). Over the course of his interactions with the experiment, ENG manufactured multiple stick tools (leafy and non) to use for honey extraction (summarized in Table 4). During his engagement with Experiment 2, ENG used the leafy stick that was provided to dip the leaves in and suck the honey from them. He also manufactured more stick tools and leaf sponges from nearby shrubs and saplings throughout the experiment to continue honey extraction.

**Figure 4:**
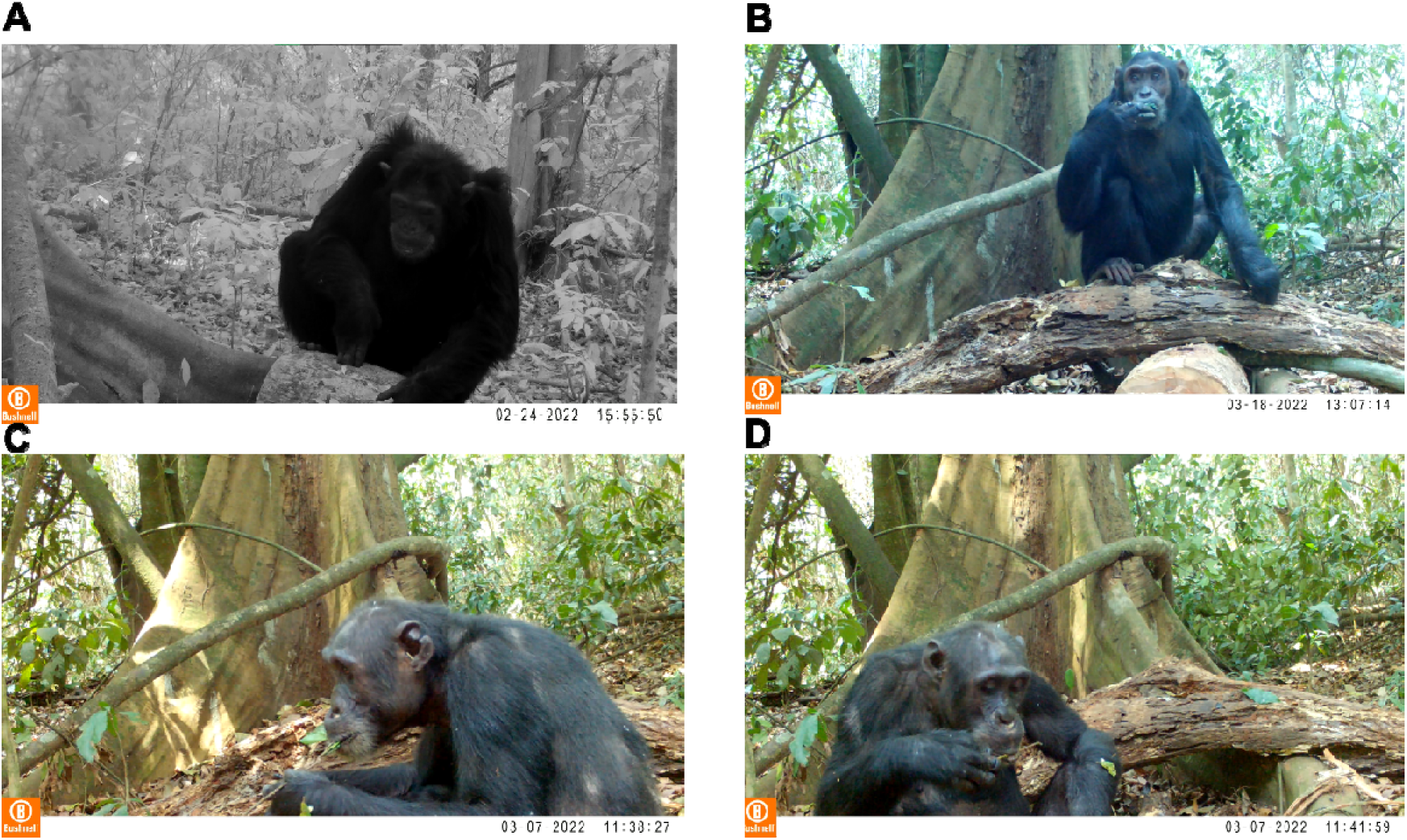
Variations in the honey-trap Experiment 1 engagement. A) An adult male using fingers to try to reach honey. B) A female chimpanzee (HON) leaf-sponging. C) An adult male (ENG) using the leafy stick as a dipping tool to extract honey. D) ENG using a bare stick to extract honey. Credits: @Kelly Ray Mannion / BPCP.

## Discussion

Field experiments have the potential to both uncover the tool behaviour of wild animals, as well as to isolate the parts of the environment that appear salient to them. Both of these goals were pursued here, as we sought to first uncover the tool use behaviour of the Mwera South chimpanzees, and second, to determine whether individuals would recognize a potential tool for the honey extraction task. In addition, our set-up in Bugoma Forest, replicating those carried out before in Budongo and Kibale forests, allowed us to directly compare results across Ugandan communities. Overall, our results in Bugoma Forest, as well as their comparison with other Ugandan chimpanzee communities illuminates our understanding of cultural dynamics between and within groups.

Our first analysis comparing the three communities of Mwera South, Kanyawara and Sonso is suggestive of differences in cultural attitudes towards a given food in chimpanzees. In particular, the Kanyawara chimpanzees appear different from the two other communities in their attitude towards honey, both in terms of engagement time, as well as by having a dedicated instrumental action to extract the honey exhibited by nearly all tested individuals. Overall, our analyses suggest that Kanyawara chimpanzees might be generally more interested in exploiting the honey when finding it compared to the Mwera South and Sonso chimpanzees. We remain cautious in our interpretations given the limited statistical power and inconsistent results across statistical methods, yet the latter should be understood in context too. Our first goal was to replicate the inter community comparison in Gruber et al. 2011, but this time across three communities. The analysis in Gruber et al. 2011 excluded two types of outliers: the chimpanzees who engaged too much with the experiment and the ones who did not engage enough. The reason for the former was to remove statistical outliers (only one individual, QT, replicated here), while the reason for the latter was conceptual: in that study, we wanted to give Sonso chimpanzees enough time for chimpanzees to use a tool, and used the median value for Kanyawara to use one (20 s) as a threshold. Over the course of the following decade, we confirmed that giving more time – or even more opportunities to use tools – to chimpanzees in Sonso would not lead them to develop more tool usage. Rather, this relationship is inverted: chimpanzees who use tools are actually the ones who consistently spend more time with the experiment extracting honey [22]. This is once again confirmed with our results in Mwera South, where tool users were amongst the individuals who engaged the longest with the experiment. A different way to interpret our results is thus that an individual who engages only 2 to 20 s with the experiment does not actually want to engage more than this, and therefore has a limited interest in the resource, as is repeatedly the case in Sonso and Mwera South. In contrast, in Kanyawara, even the low-engaging individuals engaged longer than 20 s with the experiment, and crucially, over 86% of them engaged with the same tool, sticks, to exploit this resource. On a side note, all data in Kanyawara were acquired in a matter of weeks, compared to months in Sonso and Mwera South, suggesting that the duration of engagement for the Kanyawara chimpanzees, all opportunities equal, would be much longer than for the two other communities. As such, we believe that our results support differences in *cultural attitudes* toward the resource, which can be considered as a separate component of general cultural knowledge, different from the instrumental cultural knowledge present in the community, as is exemplified by our results in Bugoma.

The results from the Bugoma honey-trap experiments support conclusions drawn from other honey-trap experiments in that chimpanzees engaging with the experiments solved it with their existing instrumental cultural knowledge, i.e. leaf-users used leaves and stick-tool users used sticks. As such, our experiments confirm rare sighting of stick usage prior to the experiments in Bugoma (see Supplemental Material and discussion below). However, our results are also unique compared to the two other communities, in that chimpanzees within a single community used two different types of tools and material (leaves and sticks) to solve the problem, a never seen before situation in over a decade of field experiments across two field sites and involving over 60 individuals. The Sonso chimpanzees only ever used leaf tools to extract the honey and did not perceive the stick tool provided to them as useful, only removing the leaves from it to sponge honey. The Kanyawara chimpanzees, in contrast, who had been documented as stick tool users prior to the honey-trap experiments [31], only used stick tools to extract the honey, foregoing using the leaves attached as sponges or for dipping, and utilizing only the bare stick part when a leafy stick tool was provided. It is important to note that all Kanyawara chimpanzees routinely use leaf-sponges for water but refrained from doing so in the honey-trap experiments leading us to conclude that the use of a tool and its contextual use can be learned separately [32]. This was seemingly the case with HON, a leaf-sponger in Mwera South who did not use any tool to extract the honey until after seeing MAM do so. A similar sequence was observed in Budongo, where PS, a male chimpanzee, engaged first with his fingers until observing RS, a female chimpanzee, leaf-sponging, and only then manufactured a leaf-sponge himself [18].

Remarkably, despite the comparatively small number of Mwera South individuals engaging in the honey-trap experiments, they exhibited all of the behaviour of individuals in previous studies including using leaf and stick tools for extraction. Additionally, we report for the first time that an individual, MAM in Mwera South, used a honeycomb to sponge the honey from the experiment. To the best of our knowledge, this has never happened in experiments conducted in other wild chimpanzee communities. As mentioned earlier, innovation is scarce and though seemingly difficult or unlikely, it does occur and has been documented in the wild before ([2] [33]). We hypothesize that a chewed honey-comb resembles a fruit wedges, which is a common product of their fruit-eating diet [34]. Nevertheless, the possibility from at least some chimpanzees to flexibly use part of their environment as tools, beyond their set function, underlines a potential larger capacity for adaptation in wild chimpanzees, as also seen in the behaviour of ENG. Indeed, this adult male was particularly inventive in the experiments and exemplifies the individual variation that can occur between chimpanzees, even from the same group, engaging in the same task. After manufacturing a stick tool on his own in Experiment 1, he manufactured both stick tools and leaf-sponges from the provided leafy stick, as well as from the experimental setting, displaying an undocumented flexibility in the honey-trap experiments so far. Crucially, while trial-and-error are expected in younger individuals while they develop their tool set, this behaviour was observed in a single adult male, for whom the cultural baggage of his community should be well acquired by that time, and possibly less flexible because of functional fixedness (see [22] for discussion). A different approach here is to acknowledge that there are clear outliers within their own communities, who are particularly fond of a given resource, and who will go to a larger extent to find creative ways to obtain it because of motivation alone. Such individuals may become innovators in their own right, although it is not clear how the two characteristics interact.

Finally, beyond the cognitive and motivational aspects we have discussed, our results stress that the ecological validity of field experiments also reveals itself in context. Indeed, the multiplication of studies, observational or experimental, of wild chimpanzees across Uganda allows drawing a larger picture of the stick tool use culture. Stick tool use happens in other Ugandan communities such as those of Bulindi, Kasongoire, Kasokwa in the Budongo-Bugoma corridor, as well as in Kibale and Kalinzu, but the evidence for stick tool use in Bugoma had been scarce since the beginning of the project. This is not unheard of as it sometimes takes decades for researchers to observe cultural behaviour in chimpanzees [35]. Prior to our experimental observations, we only had admittedly weak and indirect evidence for stick tool use. Several marks at claypits or termite mounts suggest that local chimpanzees may use sticks for digging although no video has been acquired so far. Since stick tool use behaviour occur in these manners at other field sites in Uganda, it is not outside the scope of possibilities that they occur in Bugoma as well. Berger et al. [36] theorized that the Bugoma chimpanzees would be likely to use stick tools based on ecological findings that Bugoma Forest appeared more similar to Kibale Forest than to Budongo Forest. This, coupled with knowing that innovation is rare and the instances of ENG using and recognizing stick tools, lead us to support this theory with experimental evidence, that is Bugoma chimpanzees likely have stick tool use in their cultural repertoire. As research progresses in Bugoma Forest, we will be able to determine exactly in which contexts stick tool use is occurring in the Mwera South community as well as in other chimpanzee communities in Bugoma Forest Reserve. Our results however suggest that the Mwera South chimpanzees do not associate stick use to honey in a cultural manner as the Kanyawara chimpanzees, being in that respect closer to the Sonso chimpanzees in their attitude towards honey. This outlines a conceptual distinction in the wild cultures of chimpanzees (cultural attitude vs instrumental knowledge) that warrants more investigation, while taking into account idiosyncratic variations of individuals connected to their motivation.

## AUTHOR CONTRIBUTIONS

Kelly Ray Mannion (KRM) and Thibaud Gruber (TG) conceived the ideas and led the writing of the manuscript. KRM conducted all field experiments with assistance from BPCP field assistants. All authors contributed critically to the drafts and gave final approval for publication.

## ACKNOWLEDGEMENTS

Kelly Ray Mannion and Thibaud Gruber were supported by a grant of the Swiss National Science Foundation (grant PCEFP1_186832 to Thibaud Gruber). We are greatly appreciative of the Bugoma Primate Conservation Project field assistants, staff and research coordinators for their help with fieldwork and Co-Director Dr. Catherine Hobaiter. We are grateful to Ugandan National Council for Science and Technology (UNCST), Uganda Wildlife Authority (UWA) and the National Forest Authority (NFA) for permissions to conduct this project in Bugoma Forest Reserve.

## CONFLICT OF INTEREST

There is no conflict of interest.

## Notes

https://github.com/ChimpanzeeResearch/bugoma_tool_use/tree/main

### Competing Interest Statement

The authors have declared no competing interest.

