## Supplementary for "Variation in cultural attitude, knowledge and individual motivational factors impact engagement and tool use in a field experiment in wild chimpanzees"

**Tool use observed in Mwera South community (all credits: Kelly Ray Mannion)**

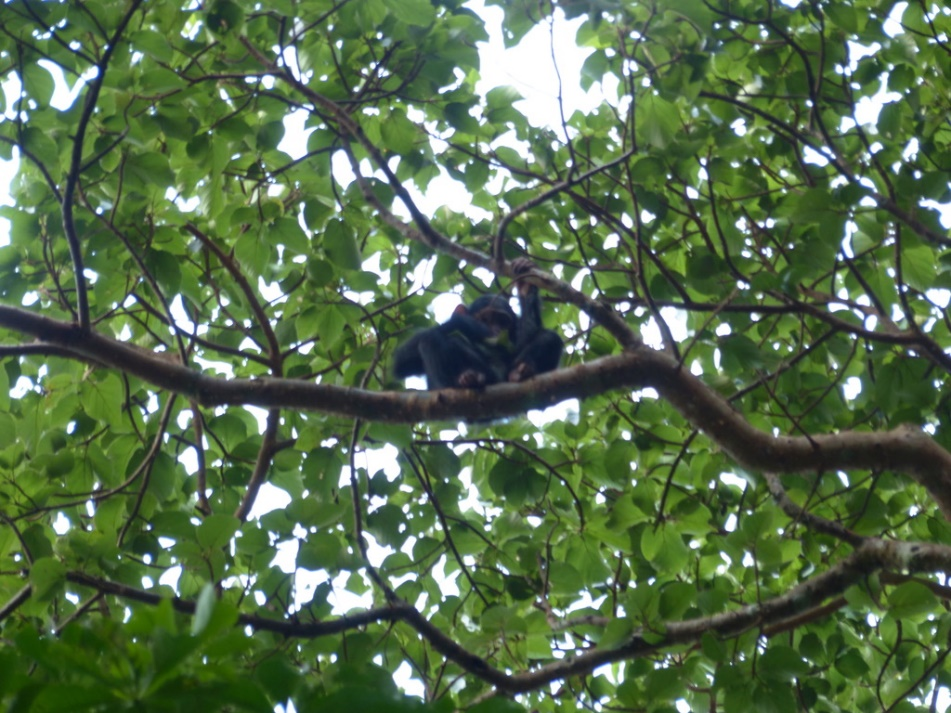

**Supplementary Photo 1:** A juvenile Mwera South chimpanzee wiping itself with leaves.

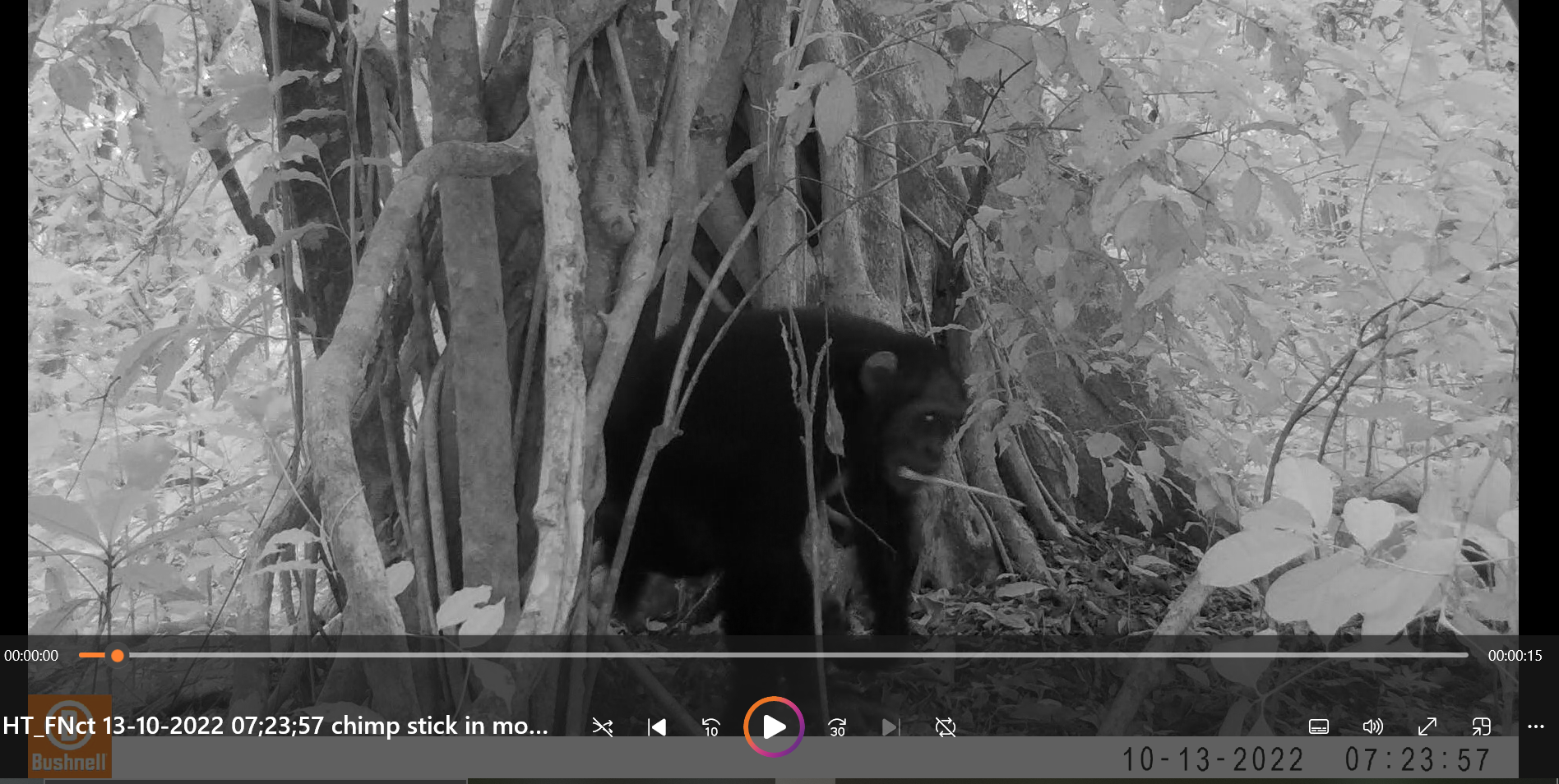

**Supplementary Photo 2:** Screenshot from a motion sensing camera trap set up in Bugoma Forest of a Mwera South chimpanzee moving with a stick in its mouth.

**Supplementary videos (all credits: Kelly Ray Mannion)**

**Supplementary Video 1:** Female Mwera South chimpanzee wiping herself with leaves. Video: [https://youtu.be/XgoXkBAJY44](https://youtu.be/XgoXkBAJY44%20)

**Supplementary Video 2:** Mwera South adult male, ENG, wiping with leaves after a copulation at the experiment location. Video: <https://youtu.be/HgcqXyZu6QA>

**Supplementary Video 3:** Mwera South adult male, MUK, leaf sponging for water from a hole in a tree. Video: <https://youtube.com/shorts/T7L5sHsm7fY?feature=share>

**Supplementary Video 4:** Mwera South adult female, MAM, using a honeycomb as a sponge to extract honey from the experiment and sharing tools. Video: <https://youtu.be/qdMIMo6kDEQ>

**Supplementary Video 5:** Mwera South female using a leaf sponge to extract honey from the experiment. Video: <https://youtu.be/fa-gd1bWqc0>

**Supplementary Video 6:** Mwera South adult male, ENG, using a leafy stick, and then a stick to extract honey in Experiment 1. Video: <https://youtu.be/fJeXbPUiXUE>
